# Inferring single-cell dynamics of a molecular reporter from unlabeled live-cell light microscopy data analyzed with delay embedding

**DOI:** 10.1101/2025.01.23.634593

**Authors:** Jalim Singh, Jeremy Copperman, Laura M. Heiser, Daniel M. Zuckerman

## Abstract

Quantification of the temporal sequence of molecular behavior in live individual cells holds promise for improving causal and mechanistic models of cell biology. In recent years, different methods for inferring molecular labeling from microscopy data have been developed, especially in the context of “virtual pathology”, but less effort has been directed to the context of single-cell dynamics and live-cell imaging. We demonstrate that phase-contrast live-cell imaging of MCF10A cells, without labeling data, is predictive of dynamical, single-cell behavior of the cell-cycle reporter Human DNA Helicase B (HDHB) – in particular, of the nuclear vs. cytoplasmic localization of this fluorescent reporter of cyclin-dependent kinase activity. Prediction quality improves substantially when temporal sequences of images are combined in a “delay embedding” framework. When different featurizations of the imaging data are examined, we find that features derived from a variational auto-encoder (VAE) outperform “classical” image features derived from shape and texture. We find the best performance, with Pearson *R∼* 0.9 on test data, using VAE features augmented by categorical predictions, all within the delay-embedding framework – in comparison to *R ∼* 0.5 based on “ordinary” regression with VAE features.

## I. INTRODUCTION

In this study, we aim to make quantitative single-cell dynamical predictions for a molecular process based on unlabeled imaging data alone. Ultimately, if extended to the omics scale, a dynamical model of molecular components – e.g., mRNA and/or proteins – would provide a resource from which numerous mechanistic hypotheses and inferences could be generated based on the temporal order of events. Omics assays of cells such as RNAseq [1–4] or proteomics [5–8] offer high-dimensional characterization of cells, but they are destructive and do not track dynamics.

It is well-appreciated that cell images contain information on molecular and microscopic aspects of cell behavior based on a variety of prior work, e.g., “virtual staining” of molecular markers from H&E images [9–11], inference of cell state and perturbation responses from images [12–16], and inference of bulk RNA levels from imaging [17]. Live-cell imaging is a non-destructive assay that can track large numbers of individual cells over time in a single experiment [18–20]. On the other hand, despite the acknowledged richness of live-imaging data [12, 17, 21–23], we are not aware of high-accuracy dynamical molecular models derived from imaging.

The present study builds directly on our prior work employing a “delay embedding” framework [24] to leverage cellular morphodynamics for characterizing cell behavior. Delay embedding improves upon conventional image analysis by directly employing the time-ordering of images. In simple terms, the “embedding” process enriches the set of features available for downstream analysis because each time point is additionally associated with a time-ordered set of preceding images. Our prior work showed that delay embedding improved the predictiveness of a Markov model of cell behavior [12] and also enabled accurate prediction of bulk mRNA levels for cells subjected to different perturbations [17]. However, the discrete morphodynamic states used in these prior studies were fairly coarse and proved inadequate for high-precision characterization of the dynamics of a single-cell molecular marker.

Our results obtained in the present study show that morphodynamical features, i.e., image features employed with delay embedding, are indeed sufficient for highly accurate dynamical single-cell characterization of the HDHB molecular marker, the human DNA Helicase B, which transitions in and out of the nucleus in correspondence with the cell cycle [25–27].

## II. METHODS

### A. MCF10A cell culture and live-cell imaging

We studied MCF10A cells, a normal but immortalized human breast epithelial cell line, treated with ligand EGF, epidermal growth factor. The EGF treatment leads to “healthy” MCF10A cells which nevertheless exhibit heterogeneity of single-cell behavior in terms of morphology, motility, and cell cycling [12, 17, 28].

Live-cell imaging was performed using three channels. Phase (gray), nuclear (red), and cell-cycle (green) reporter channels were collected at 15-minute intervals for 48 hours for four fields of view (FOVs) in each well of a plate. More information about the cell culture and imaging is given in Refs. [12, 17].

### B. Image processing and single-cell trajectory generation

Phase and nuclear channel images were segmented using the Cellpose 2.0 package [29]. A custom model built from the pre-trained *cyto* model in the Cellpose 2.0 package, was trained to segment the cytoplasm of cells by selecting images at different time points and FOVs. This trained model was used on all the images segmenting cytoplasm of millions of cells.

To get single-cell trajectories, cells were tracked across time frames using Bayesian Tracker (btrack) — a Python library [30]. In turn, these trajectories were used to prepare short trajectory “snippets” consisting of a specified number of snapshots in a sliding window manner [12].

### C. Quantification of cell-cycle via nuclear-to-cytoplasmic ratio of the HDHB reporter

Cells are sensitive to growth factors during late G2 and G1 cell-cycle phases, not during the S-phase [31]. We quantified the signal intensity of human DNA helicase B (HDHB), an established cell-cycle reporter, for every snapshot recorded in live-cell imaging [26], i.e., every 15 minutes. We utilized the segmented phase and nuclear channel images to isolate the nuclear boundary from the cytoplasm and to calculate HDHB reporter signal intensity within the nucleus and cytoplasm. In all analysis reported here, the nuclear-to-cytoplasmic (N/C) ratio is calculated as the ratio of the mean signal intensity within the nucleus and the cytoplasm.

### D. Initial featurization for snapshots

Images at single time points (snapshots) are featurized in two ways in this study. First, “classical” image features (total 95) used in this study are computed based on cell shape, size, texture, and cell-cell interactions [12]. In our prior work, these features were employed to characterize multiple cell states and and further to deconvolute bulk RNAseq data for MCF10A cells undergoing different ligand treatments [12, 17].

We also examined more modern features generated from machine learning, specifically from a variational auto-encoder (VAE). We used the Adam optimizer [32] to optimize the VAE objective function, where we set the learning rate at 0.001 [33]. The VAE is trained for 100 epochs with a batch size of 128 to extract 256 features from the live-cell images (snapshots).

### E. Delay embedding: Use of trajectory snippets

We employed delay embedding – essentially, the concatenation of time-ordered features – to enrich the feature set available for regression. Procedurally, we first featurize snapshots and then multi-snapshot “snippets.” For snapshots, we have an initial number of features *n*, either *n* = 95 classical features or *n* = 256 VAE-derived features. To implement delay embedding, we concatenated features from the preceding time-points, depending on the length of snippets *l*. Thus the total number of features for a trajectory snippet would be *n·l*: for example, a snippet length of 2 would lead to 95*·*2 classical features. However, to control computational cost, if a snippet’s total number of features exceeded 256, we applied principal component analysis (PCA) from scikit-learn to reduce it to 256. Thus, the maximum number of features employed for regression for any snippet is 256.

### F. Regression models, training data, and auxiliary features

We employed two types of regression model to predict cell-cycle information directly from the morphodynamic cell features as a regression task, namely least absolute shrinkage and selection operator (LASSO) with cross-validation [34]and the Random forest [35] as implemented in scikit-learn.

To optimize the performance of regression models, we subsampled training data so that N/C values were distributed fairly uniformly over the full dynamic range. In the raw training (and test) data, the distribution of N/C ratios exhibits two major peaks in the middle of the N/C range (not shown) presumed to correspond to different cell-cycle states. We found that using all the highly non-uniform training data led to poor regression models, essentially because models learned only the dominant, mid-range N/C ratios. To mitigate this issue, we binned our data along the full N/C range and subsampled data with N/C ratios in over-represented bins, i.e., eliminating training data from highly occupied bins, ensuring approximately equal representation across bins. This preprocessing step was applied before training of all models presented in this study. No such data subsampling was performed on test data, i.e., all trajectories and images were used in testing.

Based on visual examination of preliminary regression results, we added additional features based on a classifier trained to recognize very small and very large values of the N/C ratio. The classification of VAE-derived features is performed by constructing three classes of the N/C ratio in the ranges of [0, 0.9], [0.9, 2.7], and [2.7, *∞*]. The ranges of the N/C ratio are defined so that the first and third classes represent relatively unambiguous instances of cell-cycle states G1/S and G2/M, respectively. However, the middle class simply contains “everything else” and thus is biologically ambiguous. A convolutional neural network (CNN) [36] was trained for the classification task, and the resulting three predicted class probabilities are used to augment the features extracted from the VAE where noted.

### G. Model evaluation

The performance of the ML models was evaluated by computing mean squared error (MSE), the Pearson’s correlation *R*, and R-squared (*R*^2^) for the regression task. For classification, the confusion matrix, F1 score, accuracy, precision, and recall were used (not shown).

Training and test data consisted of separate biological replicates. Live-cell imaging was performed for two biological replicates of MCF10A cells treated with EGF, denoted EGF1 and EGF2. Each biological replicate consisted of four technical replicates in separate wells on the plate. In the present study, we trained our model on the wells of EGF1 and tested on the wells of EGF2.

### H. Implementation and Code Availability

All models were implemented using Python and the scikit-learn library. We used TensorFlow to implement the VAE model for the regression task to predict cell-cycle dynamics. The codes used in this study can be accessed at https://github.com/ZuckermanLab/CellTrajAnl.git.

## III. RESULTS AND DISCUSSION

### A. Comparison of regression performance among featurizations and models from snapshot data

As a first task, we performed regression based solely on snapshot data, i.e., standard single images of cells. Using features from the phase-contrast channel only, we trained regression models to predict the N/C ratio of the HDHB cell-cycle reporter.

Our results for snapshot-based regression (Fig. 2) show that performance can be improved substantially based on featurization and the regression model. The Classical LASSO model, utilizing traditional image features, achieves moderate correlation (R: 0.510) and reasonable mean squared error (MSE: 0.140) in predicting this ratio (see Fig. 2(a)). The VAE LASSO model, shown in Fig. 2(b), leveraging features extracted from the VAE, shows slighltly improved correlation (R: 0.531) and lower MSE (0.126), indicating a further improvement in the prediction of cell-cycle state. Fig. 2(c) shows that the Random Forest model, utilizing VAE features, demonstrates an additional small improvement in correlation (R: 0.539) and low MSE (0.124), further highlighting the importance of VAE features in predicting the N/C ratio.

**Figure 1.**
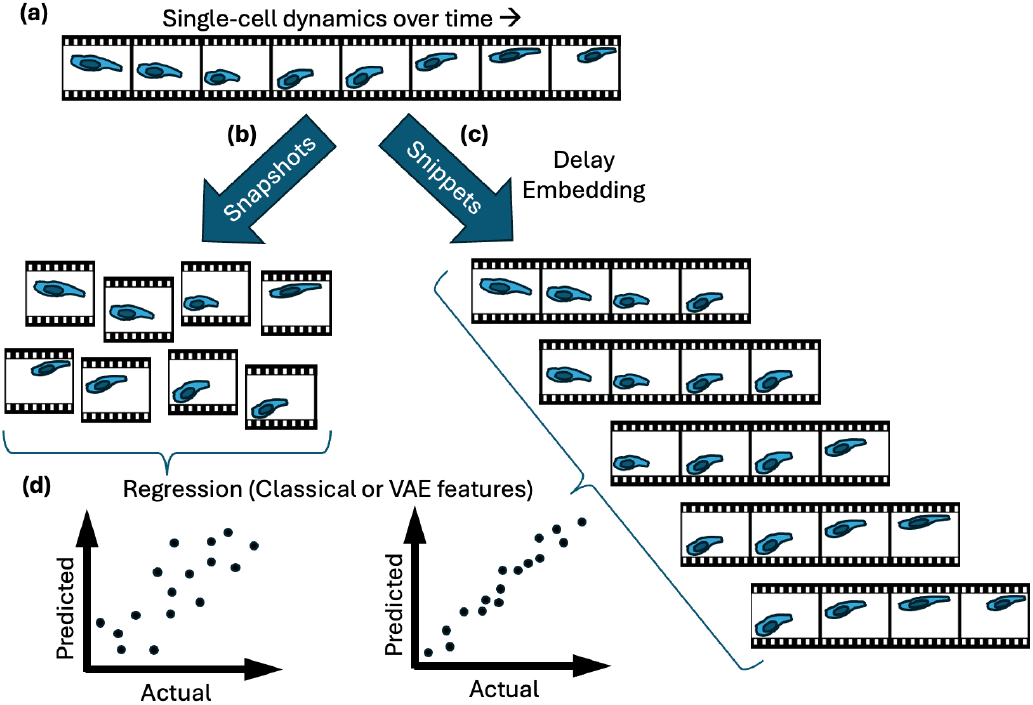
Delay embedding for machine learning of labeled features from live-cell imaging. (a) Live-cell imaging is performed and single cells are tracked over time. Cell images can be organized as (b) simple snapshots or (c) trajectory snippets consisting of multiple consecutive frames in a “delay embedding” framework. (d) Features can be extracted from the imaging data, such as “classical” or variational auto-encoder (VAE) features and then used for machine-learned regression. The present study uses only bright-field (phase-contrast) features and builds regression models for a cell-cycle reporter quantified in separate imaging channels.

**Figure 2.**
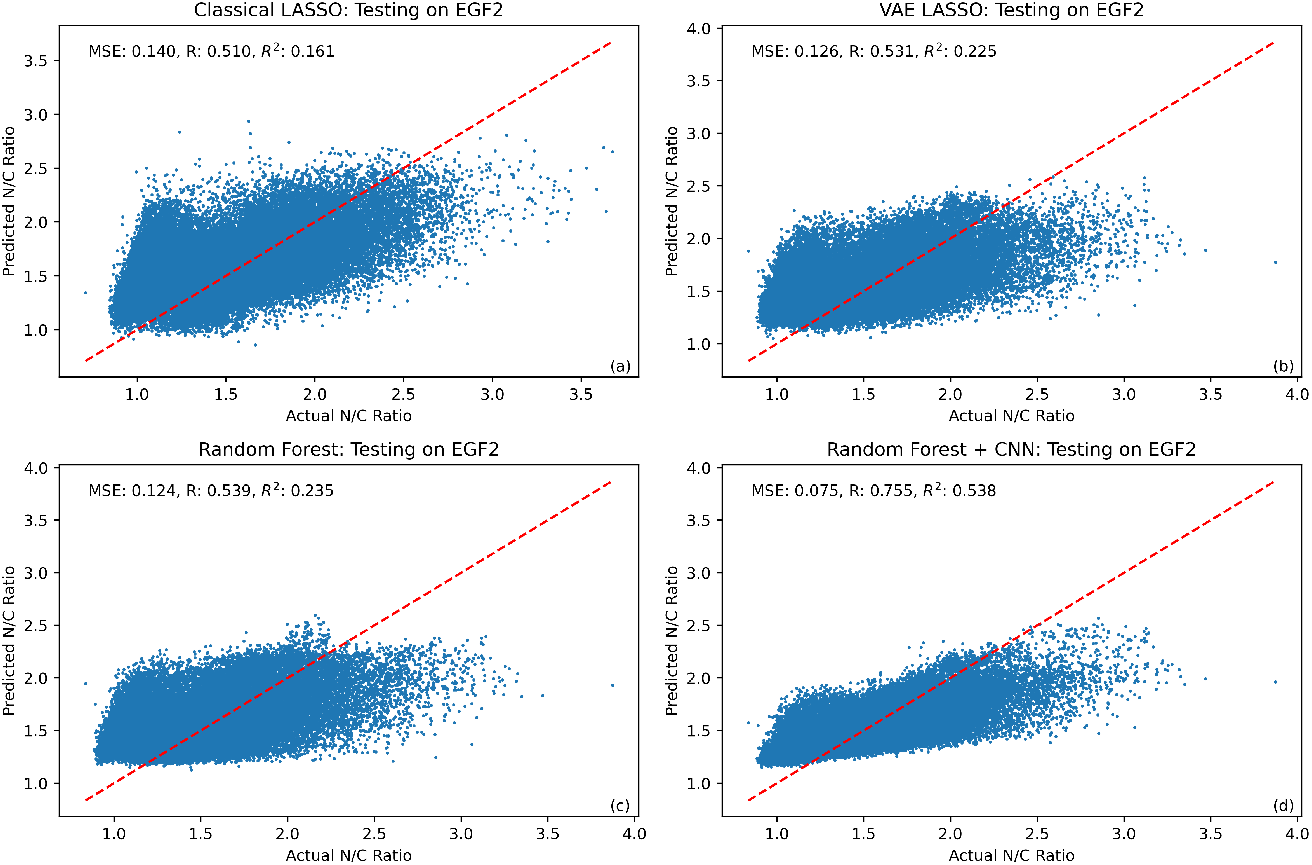
Performance comparison among ML regression models and featurizations from snapshot data. Scatter plots are shown for several models, in which each blue dot represents the predicted and actual N/C ratios for a single cell at a single time point in EGF-treated MCF10A cells. (a) LASSO on classical image features (b) LASSO on features derived from VAE, (c) Random Forest on VAE features, and (d) Random Forest on VAE features augmented from the convolutional neural network (CNN). Representative test data is shown.

Although the preceding improvements are modest, a more substantial gain for snapshot-based regression is seen when the CNN-predicted class probabilities are used in addition to the VAE features. The VAE-derived features, combined with CNN prediction probabilities, yield the highest correlation by a notable margin (R: 0.755) and lowest MSE (0.075), suggesting a robust and accurate model to predict cell-cycle dynamics (see Fig. 2(d)).

### B. Regression based on delay embedding with VAE features

We used delay embedding in an effort to improve the performance of the ML regression models. Previously, delay embedding was shown to enhance the predictability of cell trajectories represented using morphodynamic states [12]. In Fig. 3, we compare the best performing snapshot-based model (Random Forest on VAE features augmented from CNN) at different delay embedding (snippet) lengths. Pearson’s R-value consistently increases and MSE decreases with the delay embedding length (Fig. 4). This suggests an accumulation of information about cell cycle dynamics while combining cellular features due to embedding.

**Figure 3.**
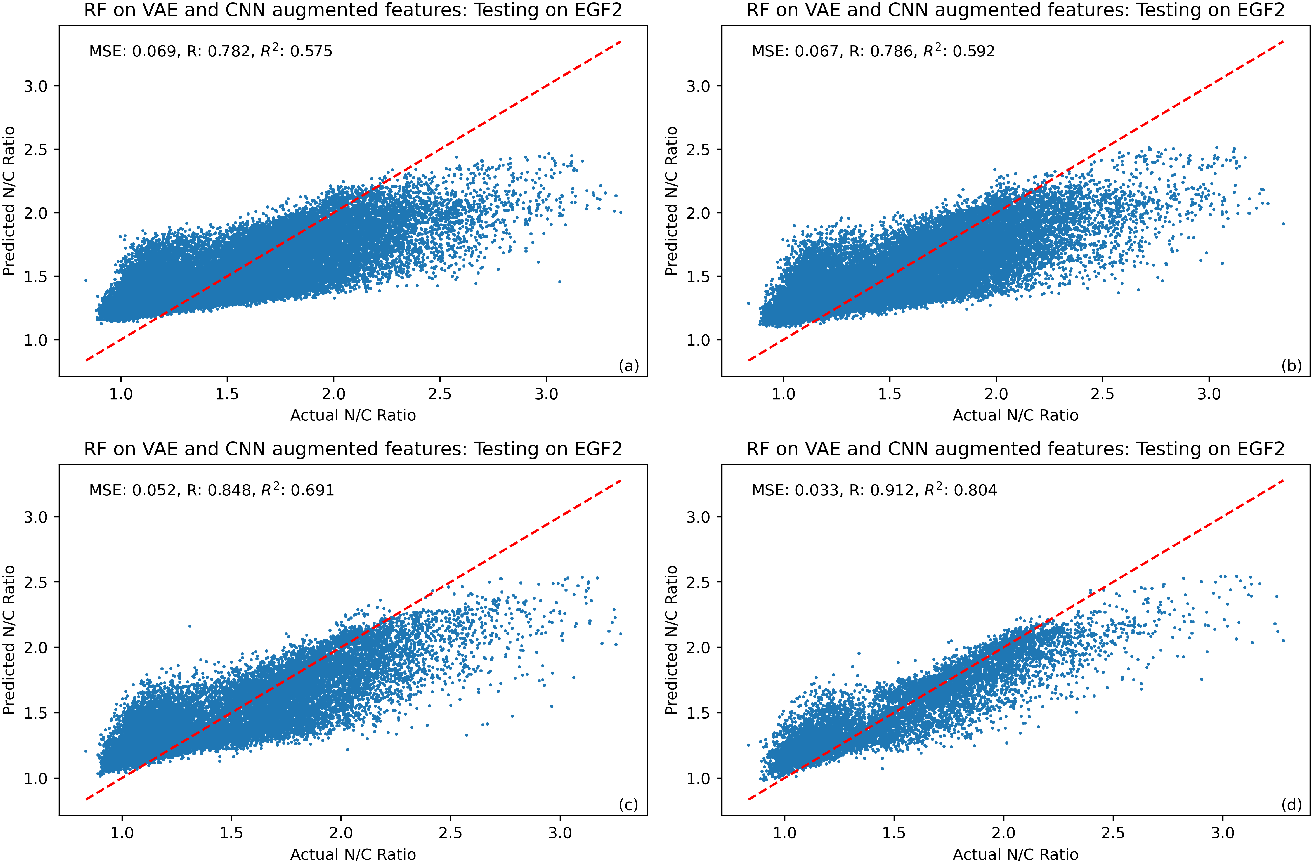
Improvement of regression performance with increased delay embedding (snippet) length. Scatter plots are shown for several models, in which each blue dot represents the predicted and actual N/C ratios for a single cell at a single time point. Plotted are Random Forest regression results based on VAE features augmented with classification probabilities from the convolutional neural network at different snippet lengths: (a) 1 hour, (b) 2 hours, (c) 5 hours, and (d) 10 hours. Representative test data is shown. Note that delay embedding employs features from multiple time points, but predictions here are made for single time points of single cells and are directly comparable to snapshot-based predictions.

**Figure 4.**
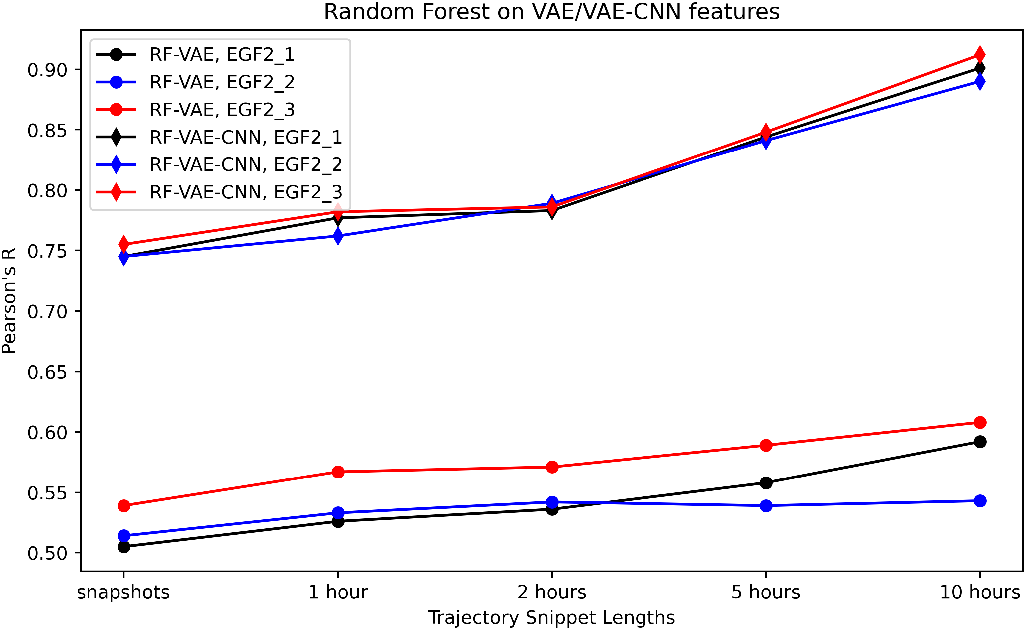
Delay embedding improves regression performance in cross-validation. Pearson’s *R* value is plotted at different trajectory snippet lengths for different test sets of EGF2 wells of the Random Forest model on VAE-derived features and the Random Forest model on VAE-derived features augmented with CNN classification probabilities.

Similar performance is found with different wells of the EGF2 replicate (data not shown).

### C. Inferred single-cell dynamics of the cell cycle reporter

As a last test, we examined model predictions for the temporal behavior of single cells. All the results considered until this point have concerned the N/C reporter ratio for static images – even when delay embedding from prior time points was used to enhance featurization.

To examine predictions of explicit single-cell dynamics visually, we show a comparison of model-predicted and measured N/C ratios over time for representative single-cell trajectories in Fig. 5. These results use the best-performing regression model determined previously, i.e., RF based on 10-hour delay-embedded VAE + CNN features.

**Figure 5.**
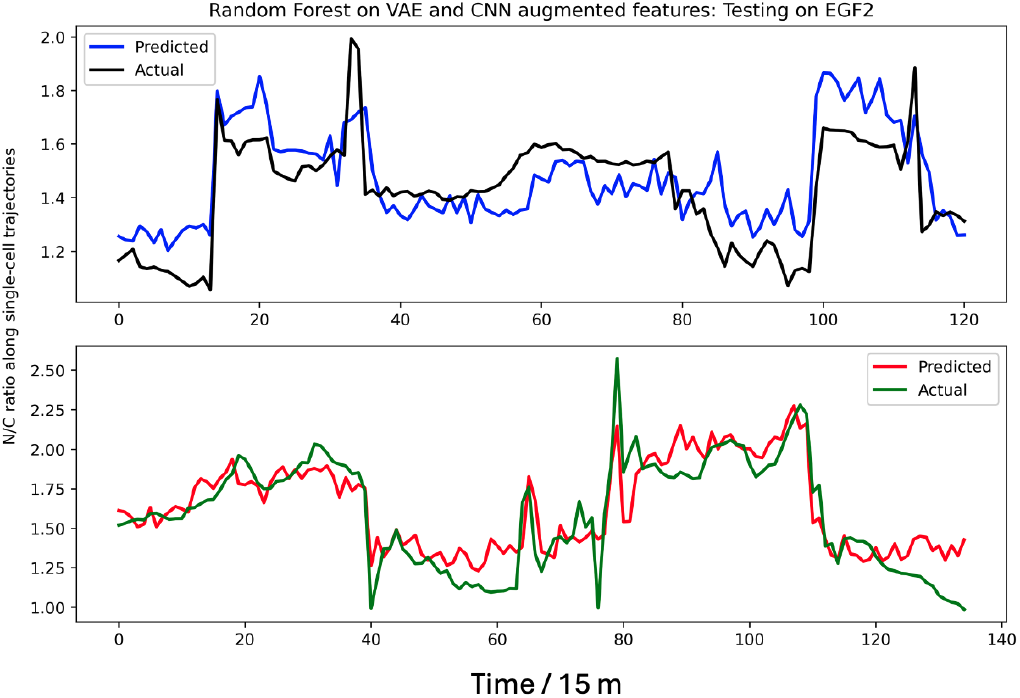
Single-cell dynamical predictions of a cell-cycle reporter. Shown are two representative single-cell trajectories of the N/C ratio for the HDHB reporter, with predictions from the Random Forest model on VAE-derived features augmented with the classification prediction probabilities.

Predicted single-cell trajectories of the of N/C ratio align closely with the actual time courses. Interestingly, the sudden jumps in the single-cell trajectories are well predicted by the model. The slower varying features are also well-predicted by our model. Note that each trajectory includes multiple snippets (Fig. 1).

We quantified the dynamical performance of our model, assessed on every available trajectory. We computed the Pearson correlation between time-matched predicted and measured values of the N/C ratio of the cell-cycle reporter. This enables examination of all trajectory correlation values, represented as histograms in Fig. 6. As expected from the static results, performance steadily improves with increasing snippet length. Although most trajectories exhibit good correlation (*R >* 0.5), some do not, which surely represents an imperfect regression model to some degree, but may also be influenced by tracking errors which would weaken the benefits of delay embedding. Nevertheless, the regression model significantly outperforms a null model based on randomly ordered predictions from the regression model.

**Figure 6.**
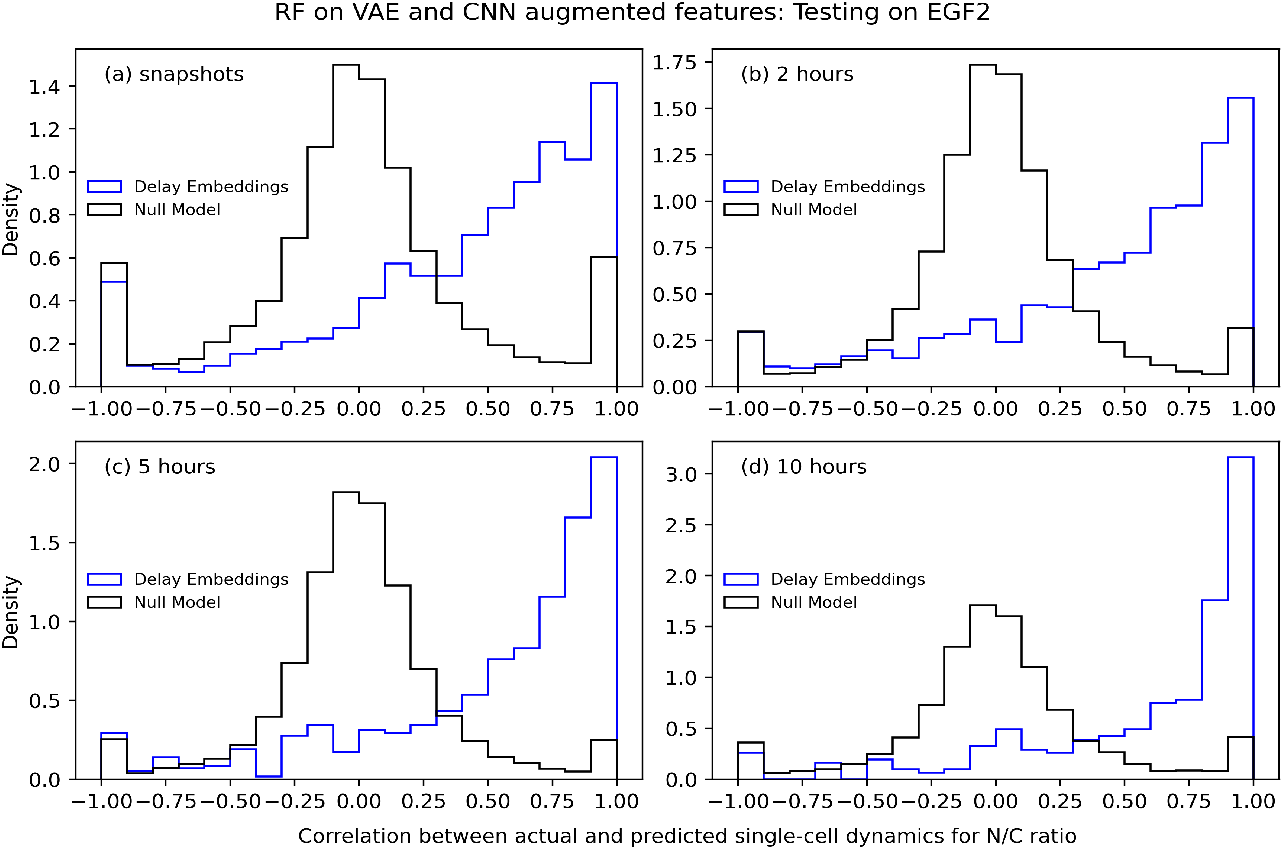
Aggregate regression performance for single-cell dynamical predictions. Histograms of Pearson *R* values are shown for time-course predictions of trajectories. Delay-embedded results (blue) represent Random Forest regression on VAE-derived features augmented with the classification prediction probabilities. The null model (black) represents the same predictions re-ordered randomly in time for each trajectory. For delay embedding, note that each trajectory consists of multiple snippets (Fig. 1).

## IV. CONCLUSIONS

The present study demonstrates that highly accurate single-cell dynamical information can be extracted from phase-contrast live-cell imaging, but that the process for doing so is not simple. Ultimately, we were able to achieve high correlation between modeled and measured values for the cell-cycle reporter HDHB in MCF10A cells. However, even sophisticated non-linear random-forest regression based on state-of-the art variational auto-encoder features did not achieve stellar performance. Top performance required the addition of several “hand crafted” features based on classifier predictions of extreme values, as well as the “delay embedding” framework. Delay embedding extracts additional information from dynamical behavior intrinsic to live-cell imaging, i.e., based on the history of cell behavior as quantified in the time course of imaging features.

## ACKNOWLEDGMENTS

The authors gratefully acknowledge financial support from the National Science Foundation (Grant MCB 2119837) and scientific input from Young Hwan Chang, Xubo Song, Mark Dane, and Ian McLean. The data supporting this study’s findings are available from the corresponding author upon reasonable request.

## References

[1] A. Mortazavi, B. A. Williams, K. McCue, L. Schaeffer, and B. Wold, Mapping and quantifying mammalian transcriptomes by RNA-seq, Nature Methods 5, 621–628 (2008).

[2] Z. Wang, M. Gerstein, and M. Snyder, RNA-seq: a revolutionary tool for transcriptomics, Nature Reviews Genetics 10, 57–63 (2009).

[3] F. Tang, C. Barbacioru, Y. Wang, E. Nordman, C. Lee, N. Xu, X. Wang, J. Bodeau, B. B. Tuch, A. Siddiqui, K. Lao, and M. A. Surani, mRNA-Seq whole-transcriptome analysis of a single cell, Nature Methods 6, 377–382 (2009).

[4] S. Anders and W. Huber, Differential expression analysis for sequence count data, Nature Precedings (2010).

[5] R. Aebersold and M. Mann, Mass spectrometry-based proteomics, Nature 422, 198–207 (2003).

[6] B. Domon and R. Aebersold, Mass spectrometry and protein analysis, Science 312, 212 (2006).

[7] S. Tyanova, T. Temu, and J. Cox, The maxquant computational platform for mass spectrometry-based shotgun proteomics, Nature Protocols 11, 2301–2319 (2016).

[8] A.-D. Brunner et al., Ultra-high sensitivity mass spectrometry quantifies single-cell proteome changes upon perturbation, Molecular Systems Biology 18, e10798 (2022).

[9] E. N. Kim, P. Z. Chen, D. Bressan, M. Tripathi, A. Miremadi, M. di Pietro, L. M. Coussens, G. J. Hannon, R. C. Fitzgerald, L. Zhuang, and Y. H. Chang, Dual-modality imaging of immunofluorescence and imaging mass cytometry for whole-slide imaging and accurate segmentation, Cell Reports Methods 3, 100595 (2023).

[10] Z. Sims, G. B. Mills, and Y. H. Chang, MIM-CyCIF: masked imaging modeling for enhancing cyclic immunofluorescence (CyCIF) with panel reduction and imputation, Communications Biology 7, 409 (2024).

[11] C. Ak et al., Multiplex imaging of localized prostate tumors reveals altered spatial organization of AR-positive cells in the microenvironment, iScience 27, 110668 (2024).

[12] J. Copperman, S. M. Gross, Y. H. Chang, L. M. Heiser, and D. M. Zuckerman, Morphodynamical cell state description via live-cell imaging trajectory embedding, Communications Biology 6, 484 (2023).

[13] G. P. Way et al., Morphology and gene expression profiling provide complementary information for mapping cell state, Cell Systems 13, 911–923 (2022).

[14] M. Haghighi, J. C. Caicedo, B. A. Cimini, A. E. Carpenter, and S. Singh, High-dimensional gene expression and morphology profiles of cells across 28,000 genetic and chemical perturbations, Nature Methods 19, 1550–1557 (2022).

[15] S. Seal, M.-A. Trapotsi, O. Spjuth, S. Singh, J. Carreras-Puigvert, N. Greene, A. Bender, and A. E. Carpenter, Cell painting: a decade of discovery and innovation in cellular imaging, Nature Methods (2024).

[16] N. Moshkov, M. Bornholdt, S. Benoit, M. Smith, C. McQuin, A. Goodman, R. A. Senft, Y. Han, M. Babadi, P. Horvath, B. A. Cimini, A. E. Carpenter, S. Singh, and J. C. Caicedo, Learning representations for image-based profiling of perturbations, Nat. Commun. 15, 1594 (2024).

[17] J. Copperman, I. C. Mclean, S. M. Gross, J. Singh, Y. H. Chang, D. M. Zuckerman, and L. M. Heiser, Single-cell morphodynamical trajectories enable prediction of gene expression accompanying cell state change, BioRxiv preprint at 10.1101/2024.01.18.576248 (2024).

[18] D. J. Stephens and V. J. Allan, Light microscopy techniques for live cell imaging, Science 300, 82 (2003).

[19] D. Kim, Y. Min, J. M. Oh, and Y.-K. Cho, AI-powered transmitted light microscopy for functional analysis of live cells, Scientific Reports 9, 18428 (2019).

[20] H. Luo, C. Jiang, Y. Wen, X. Wang, F. Wang, L. Liu, and H. Yu, Correlative super-resolution bright-field and fluorescence imaging by microsphere assisted microscopy, Nanoscale 16, 1703 (2024).

[21] S. M. Rafelski and J. A. Theriot, Crawling toward a unified model of cell motility: Spatial and temporal regulation of actin dynamics, Ann. Rev. Biochem. 73, 209 (2004).

[22] S. M. Rafelski and J. A. Theriot, Establishing a conceptual framework for holistic cell states and state transitions, Cell 187, 2633 (2024).

[23] W. Wang, K. Ni, D. Poe, and J. Xing, Transiently increased coordination in gene regulation during cell phenotypic transitions, PRX Life 2, 043009 (2024).

[24] F. Takens, Detecting strange attractors in turbulence, in Dynamical Systems and Turbulence, Warwick 1980, edited by D. Rand and L.-S. Young (Springer Berlin Heidelberg, Berlin, Heidelberg, 1981) pp. 366–381.

[25] J. Gu, X. Xia, P. Yan, H. Liu, V. N. Podust, A. B. Reynolds, and E. Fanning, Cell cycle-dependent regulation of a human DNA helicase that localizes in DNA damage foci, Molecular Biology of the Cell 15, 320–3332 (2004).

[26] S. M. Gross, F. Mohammadi, C. Sanchez-Aguila, P. J. Zhan, T. A. Liby, M. A. Dane, A. S. Meyer, and L. M. Heiser, Analysis and modeling of cancer drug responses using cell cycle phase-specific rate effects, Nat. Commun. 14, 3450 (2023).

[27] A. G. Zuniga, T. J. Aikin, C. McKenney, Y. Lendner, A. Phung, P. W. Hook, A. Meltzer, W. Timp, and S. Regot, Sustained ERK signaling promotes G2 cell cycle exit and primes cells for whole-genome duplication, Developmental Cell 59, 1724 (2024).

[28] S. M. Gross et al., A multi-omic analysis of MCF10A cells provides a resource for integrative assessment of ligand-mediated molecular and phenotypic responses, Communications Biology 5, 1066 (2022).

[29] M. Pachitariu and C. Stringer, Cellpose 2.0: how to train your own model, Nature Methods 19, 1634–1641 (2022).

[30] K. Ulicna, G. Vallardi, G. Charras, and A. R. Lowe, Automated deep lineage tree analysis using a bayesian single cell tracking approach, Frontiers in Computer Science 3, 92 (2021).

[31] S. L. Spencer, S. D. Cappell, F.-C. Tsai, K. W. Overton, C. L. Wang, and T. Meyer, The proliferation-quiescence decision is controlled by a bifurcation in CDK2 activity at mitotic exit, Cell 155, 369–383 (2013).

[32] D. P. Kingma and J. Ba, Adam: A method for stochastic optimization, arXiv preprint at Adam optimizer (2014).

[33] J. Burgess, J. J. Nirschl, M.-C. Zanellati, A. Lozano, S. Cohen, and S. Yeung-Levy, Orientation-invariant autoencoders learn robust representations for shape profiling of cells and organelles, Nat. Commun. 15, 1022 (2024).

[34] R. Tibshirani, Regression shrinkage and selection via the lasso, Journal of the Royal Statistical Society: Series B 58, 267–288 (1996).

[35] L. Breiman, Random forests, Machine Learning 45, 5–32 (2001).

[36] Y. LeCun, Y. Bengio, and G. Hinton, Deep learning, Nature 521, 436–444 (2015).

